# Full-length αIIbβ3 CryoEM structure reveals intact integrin initiate-activation intrinsic architecture

**DOI:** 10.1101/2022.12.19.520944

**Authors:** Tong Huo, Hongjiang Wu, Zeinab Moussa, Mehmet Sen, Valerie Dalton, Zhao Wang

## Abstract

Integrin αIIbβ3 is the key receptor regulating platelet retraction and accumulation, thus pivotal for hemostasis, and arterial thrombosis as well as a proven drug-target for antithrombotic therapies. Here we resolve the cryoEM structures of the intact full-length αIIbβ3, which covers three distinct states along the activation pathway. Here, we resolve intact αIIbβ3 structure at 3Å resolution, revealing the overall topology of the heterodimer with the transmembrane (TM) helices and the head region ligand-binding domain tucked in a specific angle proximity to the TM region. In response to the addition of a Mn^2+^ agonist, we resolved two coexisting states, “intermediate” and “pre-active”. Our structures show conformational changes of the intact αIIbβ3 activating trajectory, as well as a unique twisting of the lower integrin legs representing an intermediate state (TM region at a twisting conformation) and a coexisting pre-active state (bent and opening in leg), which is required for inducing the transitioning platelets to accumulate. Our structure provides for the first time direct structural evidence for the lower legs’ involvement in full-length integrin activation mechanisms. Additionally, our structure offers a new strategy to target the αIIbβ3 lower leg allosterically instead of modulating the affinity of the αIIbβ3 head region.

## Introduction

Platelets are fundamental to preventing hemorrhaging at sites of vascular injuries through thrombosis, a healthy response to injury intended to stop and prevent further bleeding. However, their functions are tightly controlled since abnormal thrombosis can cause life-threatening health problems, when a clot obstructs blood flow through healthy blood vessels in the circulatory system.^1^ lntegrins are a major family of cell surface receptors which platelets use to surveil their environment and are essential to platelet activation.^2–6^ lntegrins are multifunctional, they mediate the adhesion of cells to the extracellular matrix and to other cells, plus they participate via intracellular and extracellular signaling in diverse cellular processes including cell growth, migration, and differentiation.^5–8^ They are obligate heterodimers composed of a pair of α and β subunits.

αIIbβ3 is the major integrin expressed on platelets.^5,6^ Both the αIIb and β3 protomers are single-pass transmembrane proteins,^8^ (also known as a bitopic protein) each with a short cytoplasmic tail and a large extracellular domain responsible for heterodimer and ligand interactions. Integrin αIIbβ3 plays an integral role in thrombosis.^9,10^ Functionally, integrin αIIbβ3 transmits bidirectional signals across the cell membrane, in both “outside-in” and “inside-out” directions.^11^ Mechanistically, either extracellular or intracellular signals lead to eventual conformational changes in αIIbβ3 to an activated state, which serves as one of the final steps in platelet activation and thus plays a pivotal role in thrombosis.^12^ αIIbβ3 direct involvement in the regulation of thrombosis makes it an appealing target for therapeutic strategy development.

Prior structural studies on integrin αIIbβ3 and its homologs have focused on individual domains namely the ectodomain and the headpiece.^13–16^ Due to the flexibility of the linkers between domains, the detailed full-length structure has remained elusive, restricting our understanding on the precise arrangement of each domain with respect to one another. The orientations between the ectodomain and the TM helices are exceptionally fundamental to fully understanding the integrin conformational changes on the surface. Such knowledge will provide crucial insights into the mechanism of ligand binding modulation and of coupling between extra- and intra-cellular domains, which allows the signal to be transduced across the membrane. Previous studies on the structural determination of integrin αIIbβ3 either were restricted to the head region or only yielded low resolution structures by electron microscopy of negatively stained protein.^16–21^ Whereas the current understanding of integrin activation process implies a massive intermediate conformational change from inactive to active states, it requires a structural understanding of these intermediate states in detail.^22^ Relative organization between regions, which in turn is regulated by individual domain conformations, defines the state of an αIIbβ3 integrin and provides a structural basis for the mobility of the whole molecule.

Integrin αIIbβ3 undergoes extensive glycosylation, which shows its necessity for dimer formation and its physiological functions.^23^ However, most structural studies have used recombinant systems to obtain integrin proteins, and not all glycosylation sites and types of glycans at each site can be retained using such systems. In this study, we report the first structure of full-length integrin αIIbβ3 from native sources, with native glycosylation. The series of structures reported here provide new insights into the mechanism of integrin activation and its role in thrombosis, providing new opportunities for structural targets for future therapeutics against platelet-linked cardiovascular diseases.

## Result

### Overall structure of native integrin αIIbβ3 in its inactive state

We solved three distinct structures of αIIbβ3 induced by different combinations of divalent cations: state I (5 mM Mg^2+^/1 mM Ca^2+^); state II (1 mM Mn^2+^/0.2 mM Ca^2+^); and state III (1mM Mn^2+^/0.2 mM Ca^2+^), with states II and III co-existing, but separable structurally from cryo-EM data. Full-length integrin αIIbβ3 was purified directly from human platelets after solubilizing in detergent (see Methods). Using single particle cryoEM, we solved the full-length native structure of integrin αIIbβ3 at 3 Å resolution including both full ectodomain and transmembrane region in state I. All domains could be clearly assigned and grouped into three large regions, which are head, leg, and TM regions, according to their local positions. The detailed architecture variations in intact integrin at different conformational states among all three regions is still unclear.

In the absence of agonist, purified integrin αIIbβ3 exhibited a bent form, indicating the inactive state (state I) based on the overall features.^8^ This conclusion is supported by the ligand binding domain located at the interface between the β-propeller domain from αIIb and β1 domain from β3 shown in Fig. 1 (see details in Methods). The head and leg region form a sharp angle (~70°) between each other, defining the orientation of the whole extracellular domain and characterizing the bent form of integrin αIIbβ3. The head and leg region form the whole ectodomain that is anchored to the membrane by the TM region. Although the ectodomain has been investigated previously, it is remarkable that in this structure, the TM region topology in the full-length protein is determined for the first time. The architectural relationship between the TM and leg regions is clarified, along with the orientation of integrin αIIbβ3 towards the membrane. In our structure, the Calf-2 domain from αIIb and tail domain from β3 position the single transmembrane helix anchoring through the detergent shell that mimics the cell membrane, and form a ~60° degree angle with it. In contrast to previous schematic models proposed in their papers,^24,25^ which depicted a vertically standing integrin with its head region facing the membrane, our structure showed αIIbβ3 adopted a tilted orientation that consequently would make the head region and the ligand binding site more accessible.

**Figure 1.**
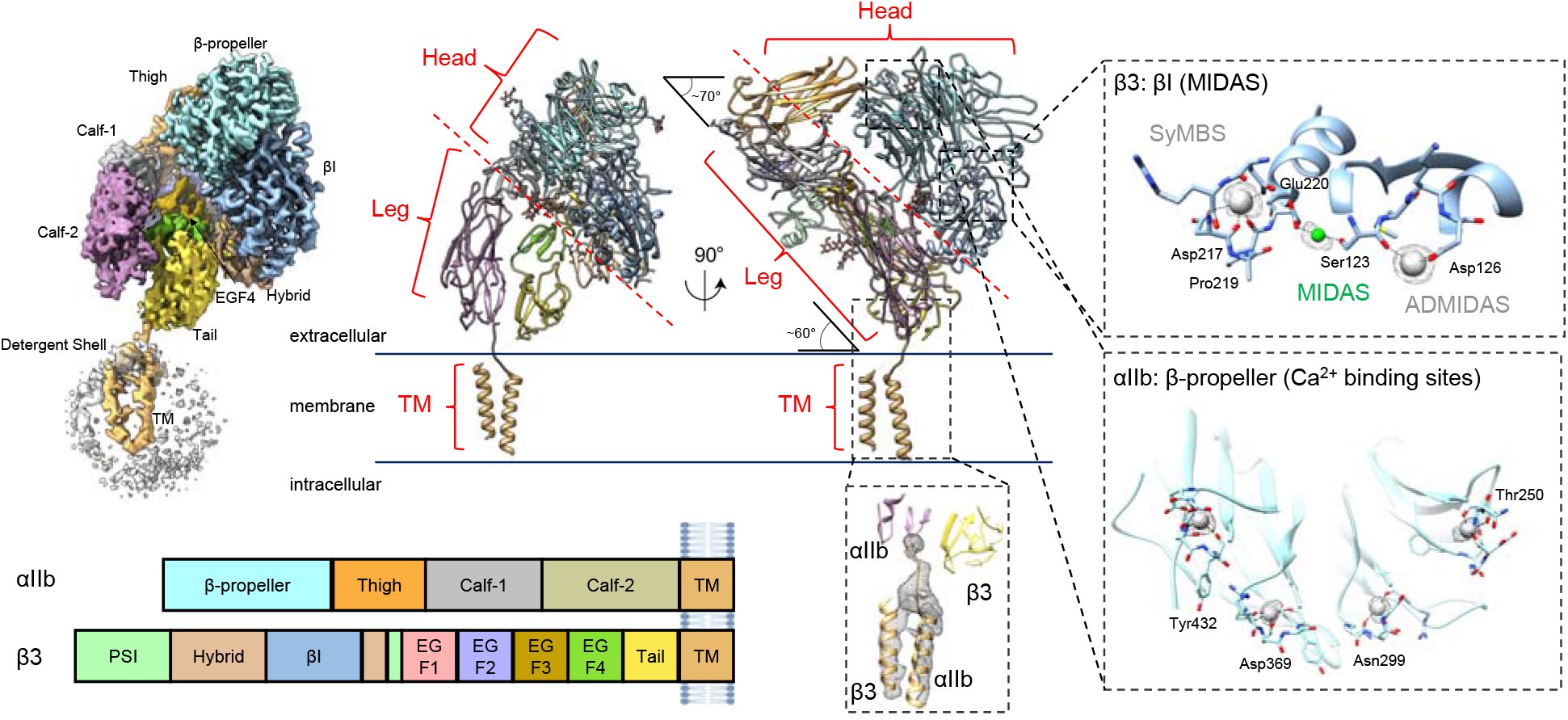
Cryo-EM single-particle reconstruction and atomic model of inactive integrin αIIbβ3. A) The structure is a heterodimer composed of αIIb and ß3 molecules. Both integrin molecules within the αIIbß3 heterdimer, αIIb and ß3, are clearly identifiable and well resolved. All five αIIb domains (ß-propeller, thigh, Calf-1, Calf-2, and TM) and all nine ß3 domains (ßI, Hybrid, PSI, EGF-1,2,3,4, Tail, and TM) are clearly assigned. B) Regions of integrin. Based on the domain organization, the whole integrin structure could also be split into three regions: Head, Leg, and TM. The integrin stands on the cell membrane in a titled orientation, with a ~70° angle between the Head and Leg regions, and ~60° between the Leg and TM regions. The inset figures show the densities for ion binding at MIDAS, ADMIDAS, SyMBS, and ß-propeller as well as a density for the TM region. C) Domain demarcation along the integrin amino acid sequence based on the structural annotation. Each domain was colored differently in the map and structure, with the same color scheme used across all figures in this paper.

The head region is critical for αIIbβ3 integrin activation, as the head region contains both the ion-binding and ligand-binding sites. The β-propeller from αIIb and the βI domain from β3 together form the main part of the integrin head region. The ion binding pockets including SyMBS (synergistic metal ion-binding site), MIDAS (metal ion-dependent adhesion site), and ADMIDAS (adjacent to MIDAS) residing in the head region are well-resolved in our structure and the coordinated ions could also be identified(Fig. 1, Supplemental Fig. 1). All three ions were resolved clearly in our structure, and there is no conformational difference compared to previous crystal structures of only the head region fragment focusing on the extracellular domain,^15^ which indicates the stability of this binding site is not likely to be impaired due to loss of the TM region. Key residues participating in ion coordination include Asp126 for ADMIDAS; Ser123 for MIDAS; and Asp217, Pro219, and Glu220 for SyMBS, all of which are mostly conserved among the RGD-binding integrins. We also observed in the distal location of the ligand-binding site for the β-propeller, four Ca^2+^ ions coordinated with adjacent residues to further stabilize the β-sheets (Fig. 1).

In the present study, the TM region was first resolved together with the connected leg regions. Although the TM region only accounts for less than 10% of integrin residues, it is a critical component of the mechanism. It undergoes movements to transduce the outside-in or inside-out signals.^26^ However, the relative orientation between the TM region and extracellular domain had yet to be determined before this study, excluding some low resolution structures.^27^ To deal with the low signal-to-noise ratio (SNR) and TM region flexibility, we employed 3D focus classification to sort out particles and resolved two helices from αIIb and β3 and the linker between Calf-2 and αIIb helix. The two helices adopted a twisted conformation and went across each other forming a knot at the membrane proximal site. The loop linker between the TM region and whole extracellular domain renders the potential flexibility for the integrin to unwind and separate the helices, which is the prerequisite of activation.

### Glycosylation sites

Aside from the TM region, our structure resolved all glycosylation sites that were uncertain in the prior ectodomain-only structure.^15^ Since our integrin αIIbβ3 was directly obtained from the membrane extraction of platelets, it likely reflects the physiological glycosylation situation. In this structure, we identified eight N-linked glycosylation sites (Fig. 2). αIIb and β3 each have four glycosylation sites, with β3 having more complicated glycosylation.

**Figure 2.**
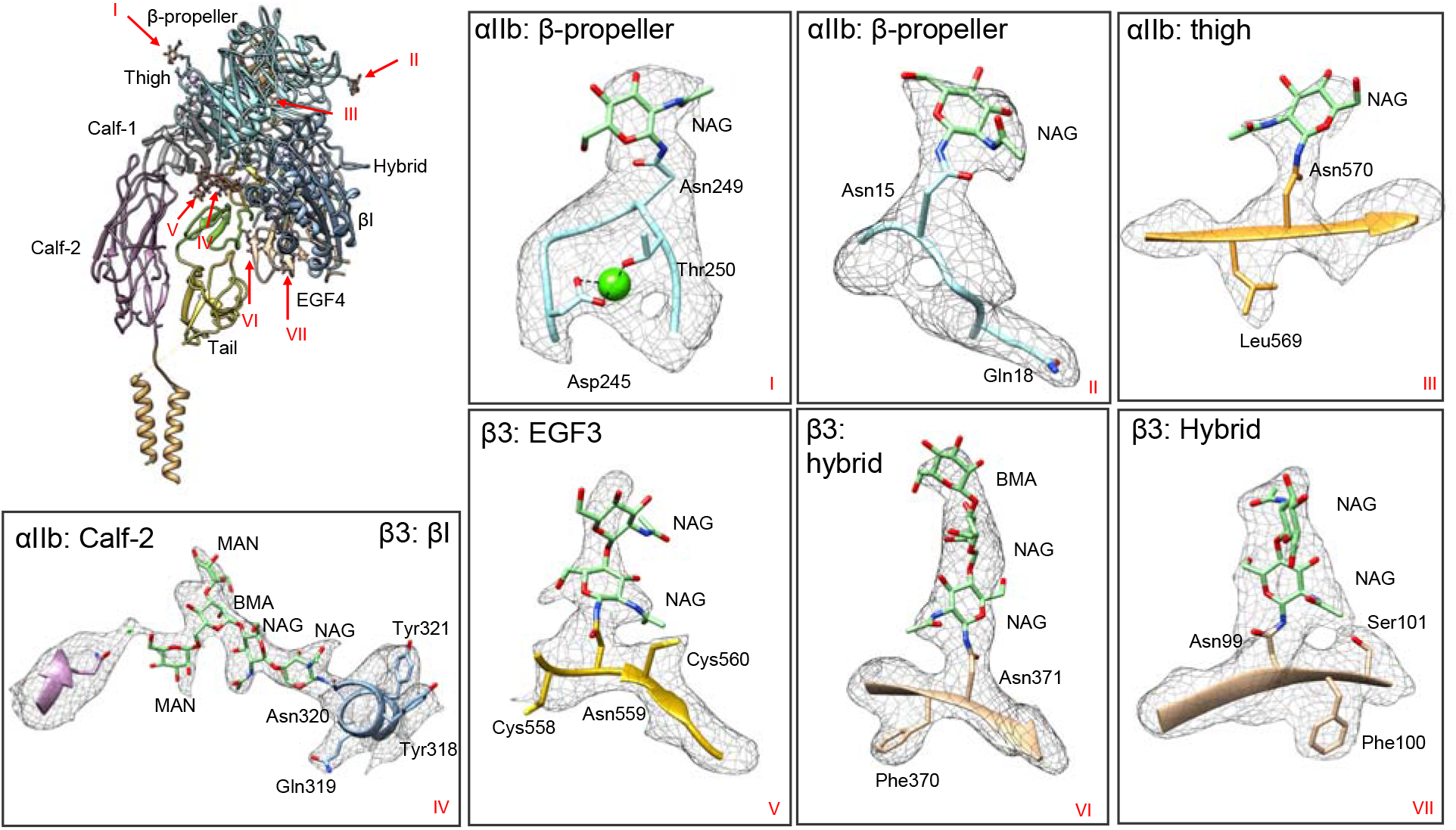
Glycosylation of integrin αIIbβ3. There are seven glycosylation sites resolved in this structure. The global distribution of the sites is shown, which reveals that most glycosylation sites are located in the Head region. The inset figures show the densities for each glycan built on the structure. We resolved this structure, for the first time, to prove that Asn931 in integrin αIIbß3 has N-linked glycans, though glycans at this site were not built into our final model due to low resolvability. At Asn320 in the ß3-ßI domain, mannose observed in this study interacts with the αIIb-Calf-2 domain through a hydrogen bond introduced by a water molecule.

In the previous αIIbβ3 structure, three N-linked glycan sites from αIIb were resolved and identified (Asn15, Asn249, and Asn570).^15^ The protein obtained from natural extract in this study also exhibited the same glycosylation pattern for these three already known glycan sites. In addition, we reveal a previously proposed but never resolved glycan site Asn931. Extra density was found connected to Asn931 indicating, for the first time, that N-linked glycans are attached to this site (Supplemental Fig. 3), though it was formerly predicted as a glycosylation site.^28^

The β3, Asn99, Asn320, Asn371, and Asn559 could be identified as N-linked glycosylation sites by the presence of the density for N-glycans, which is consistent with previous structural studies. However, N-linked glycosylation site Asn320 (β3) was identified to have five saccharide residues compared to the two saccharide residues found in the previous study.^15^ Since the glycosylation tree is located at the interface between αIIb and β3, the protruding glycans establish interactions between αIIb and β3. By introducing a water molecule, the mannose interacts with Gln821 from the αIIb Calf-2 domain through hydrogen bonds, which tethers the head to the leg and stabilizes the bent conformation (Fig. 2 IV). In addition, the dimerization of αIIb and β3, which is the prerequisite to becoming a functional unit, is reinforced by this interaction. However, since the interaction is regulated by the hydrogen bonds donated by water, the interaction is transient and easily disturbed, providing the potential for conformational changes between states. The structural details of interaction between αIIb and β3 introduced by glycans are observed for the first time in our structure, and they reveal the importance of glycosylation in the activation of αIIbβ3 as well as other homologous integrins.

### Transition from inactive to an intermediate state

The replacement of Mg^2+^ with Mn^2+^ mimics the inside-out signal transduction and triggers the activation of αIIbβ3, as deduced from recognition of Mn^2+^ induced conformation-specific antibodies.^29^ We replaced Mg^2+^ with Mn^2+^ during the purification and determined the αIIbβ3 structure bound with Mn^2+^. We found two new conformations with a particle ratio of 1:1, exhibiting domain movement and a partially opening. The structures were determined at 3.12 Å (hereafter referred to as the “intermediate state”) and 5.46 Å (referred to as the “pre-active state”) resolutions respectively. Previous structural studies of an Mn^2+^ bound integrin, which were limited to a negative stained TEM result, showed that the integrin may exhibit both bent and open state, but provided no additional high resolution details.^30^ Our cryoEM study did not resolve the integrin in a fully open and typically active state, which was inferred by previous studies on the isolated head region.^31^

Since this new structure cannot be assigned to any known state, we name it an “intermediate state” in this research. At 3.12 Å resolution, substantial conformational changes between the intermediate and inactive state of αIIbβ3 were found between these two structures. Readily discernible conformational changes and domain movements were found, even though both still adopt a bent orientation. To better elaborate the allosteric changes among all states, β-propeller domains from inactive and intermediate states were superimposed and resultant relative orientations of other domains between the two states were examined (Fig. 3). Big movements mainly appear at the leg region including the Calf-1 (α-Calf-1) and Calf-2 (α-Calf-2) domains from αIIb, and the EGF-4 (β-EGF-4) and Tail (β-Tail) domain from β3 indicated by the RMSD values for the domains ranging from 2.7 to 8.4 Å (Fig. 3).

**Figure 3.**
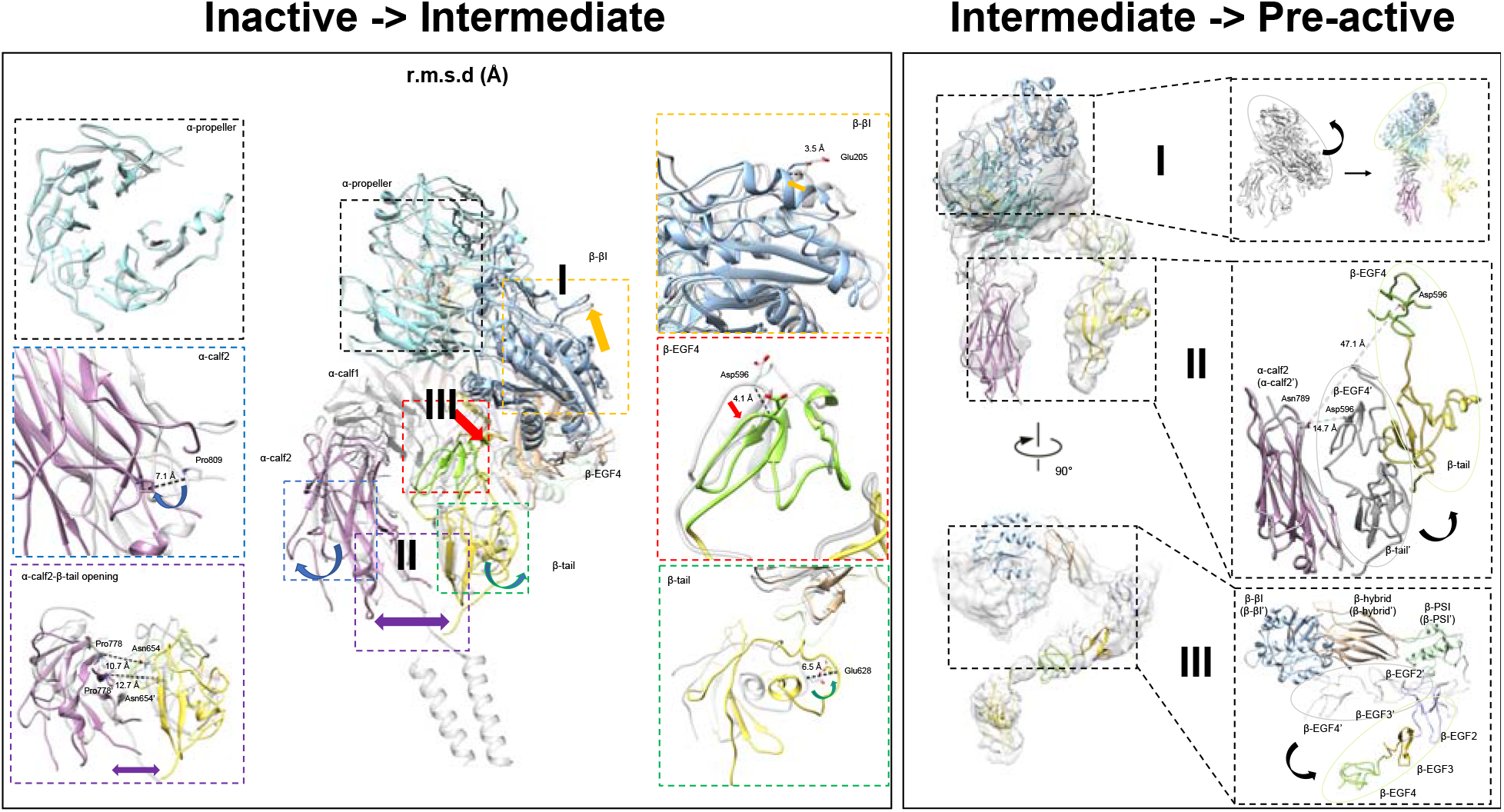
Transition from inactive state to intermediate state and pre-active state. A) Domain shifts between the inactive state and intermediate state. The structure in the intermediate state is marked with colors, while the inactive state is gray. The ß-propeller domain is superimposed to investigate shifts occurring on other domains, and C-alpha RMSD for each pair of domains is reported in the inset table. Regions I, II, and III recapitulate the movements the structure undergoes. B)Domain movement between the intermediate state and pre-active state. The structure and label for the pre-active state is marked with colors, while the intermediate state is gray. At regions I and II, α-Calf-2 includes two superimposed states, which shows a turning head region and simultaneous leg region separation. At region III, ßI and the hybrid domains in ß3 are superimposed to demonstrate the ß3 leg region swinging away.

The movements appeared mainly in three regions (**I**, **II**, and **III** in Fig. 3), as if there were three forces from different directions affecting the αIIbβ3 heterodimer. The region **I** showed a potential turn of the head region represented by a 3.5 Å shift in backbone at Glu205 from the βI domain, indicating the head region in the β3 molecule underwent a lift-up to facilitate the separation from the leg region. The region **II** has a split effect at the leg region, which is pulling apart the α-Calf-2 and β-Tail domain. Both the α-Calf-2 and β-Tail swing outwards separating the two legs, which is exemplified by the distance between α-Pro778 and β-Asn654 changing from 9.5 Å to 11.5 Å. The region **III**, mainly located at the β-EGF-4 domain, generated a shift moving away from the β3 head region as well as αIIb leg region. In summary, when integrin αIIbβ3 was treated with Mn^2+^/Ca^2+^ instead of Mg^2+^/Ca^2+^, αIIbβ3 exhibited domain movement shown as a head region lift-up and leg region separation and thus a potential to open. All three kinds of movements happened simultaneously to potentially open the αIIbβ3. Region **I**, **II**, and **III** exhibited the potential to turn the head region, to separate the leg region, and to break the connection between the head and leg regions, respectively. Consequently, TM helices were also separated by the aforementioned movement, making the signal too weak to resolve the individual transmembrane helix in our final structure of this state.

### A pre-active form captured after the intermediate form

Given the conformation of the Mn^2+^ bound integrin, αIIbβ3 is highly dynamic, as shown previously^32^ as well as by this study, we demonstrate both conformations coexist simultaneously in the same ratio. In our study, in addition to the intermediate form of αIIbβ3, another new form was also found during data processing, providing a new view of the molecular mechanism underlying the activation of αIIbβ3 (Fig. 3) in presence of Mn^2+^.^29^ Compared to the intermediate state, domain movement and even rearrangement continued to occur at regions **I**, **II**, and **III**, indicating this is a resultant or subsequent state after the intermediate state. We named this state the “pre-active” state.

When α-Calf-2 domains from intermediate and pre-active states are superimposed, variations at region **I** in the head region are the most obvious movement if viewed in the front (Fig 3). As the β-βI domain continues to lift up, generated from the intermediate state, it forms a conformation that significantly turns in the counterclockwise direction, which makes the head region move farther away from the membrane. At region **II**, the legs were separated wider than (directly measured Asn789 (α-Calf-2)-Asp596 (β-EGF4) residues distance, changed from 14.7 Å to 47.1 Å between the two states) all other states, and interactions between the α-Calf-2 and β-Tail domain could have already been disrupted at this distance. Although the density of the TM region could not be seen in this state, it is reasonable that the two TM regions from αIIb and β3 could also rotate and disjoin from each other in the scenario of a leg region separation given the distance. In addition, separation between the head and leg region also became more obvious in region **III**. The leg region, including β-EGF-2, 3, and 4, swung away from and thus formed a bigger angle with the head region. The loss of interaction between the head and leg domains, caused by the large movement, unleashed the restrictions that existed in the inactive state and provided flexibility for the integrin to be regulated. In summary, the pre-active state exhibited a subsequent conformation of the intermediate state reflected by the continuous movement occurring at regions **I**, **II**, and **III**. The resultant effect made the integrin αIIbβ3 undergo both intra-molecule separation (head and leg domain in β3) and inter-molecule separation (leg regions from αIIb and β3).

## Discussion

In this study, we resolved the structures of three integrin αIIbβ3 states in the presence of Mg^2+^ or Mn^2+^, revealing the conformational changes between different states, which shed light on the underlying mechanism of activation. We extracted the integrin αIIbβ3 directly from platelets to keep the structure close to the physiological state, especially for the glycosylation.^33–35^

Structures for both the extracellular domain^16^ and isolated TM region^34^ have been reported previously. However, the spatial relationship between the two parts has not been described. In our structure, we demonstrated that the TM region forms a 60° angle with the leg region, which means the leg region is not perfectly vertical against the membrane as previously proposed.^21,36–38^ The coiled loop linking the extracellular domain and TM region renders the flexibility of the integrin so that the extracellular and intracellular domain could have an independent conformational change, which provides the structural basis for bidirectional signaling. On the other hand, the orientation adopted by the integrin also lifts the head region up, which shows the distance between the head region and cell surface would be larger. In our model, the distance from membrane to the head region is around 5.25 nm compared to 2.41 nm showed in previous suggested models.^35,39^ (Supplemental Fig. 4) Since the head region, where the RGD-motif resides, is important in terms of ligand-binding, the longer distance to the platelet membrane would make the head region more accessible for the large population of integrin ligand proteins which contain the “RGD” and “KQADV” motif but do not share similar overall protein architecture. It is also noteworthy that the resolvability of the TM region of integrin αIIbβ3 could shed light on other bitopic proteins.

As a cell membrane glycoprotein, integrin αIIbβ3 carries several N-glycans on the extracellular domain. However, it remains unclear how the glycosylation works, it is tempting to speculate the N-glycans would be involved in the structural changes and thus the activation of integrins.^28^ Without predictions or partially resolved,^15,28,31,40,41^ our structure resolves eight glycosylation sites in the intact integrin. Among these sites, Asn931 is located at the leg region and closer to the cell membrane compared with the other glycosylation sites within αIIb, it is likely that our structure study revealed initial conformational changes by leg region movement which may play an important role during the activation process. A previous mutagenesis study on the glycosylation sites in αIIb has also demonstrated that, among all Asn substitutions in the αIIb subunit, only a N931Q mutation exhibited a decrease in both activation response and protein expression of β3.^28^ For the glycosylation at Asn320 from β3, previously resolved structures either identified only two glycans in an ectodomain structure or five glycans in a head region only structure. Thus, it is hard to clearly demonstrate how the glycosylation of Asn320 participates in the activation of the integrin. In contrast, in our full-length structure, there are five glycan residues that could be seen attached to Asn320. Asn320 was previously proposed to block the leg-proximal end of the ligand binding site in the αIIbβ3 integrin.^28^ This study reveals a longer glycosylation tree, and it shows the core glycosylation tree pointing at the Calf-2 domain (Fig. 2). It is highly conceivable that Asn320 would be more likely to play a supplemental role to connect αIIb and β3 via the glycans instead of regulating the ligand bind. In this structure, a water molecule was found to help establish the interaction between αIIb and β3 and stabilize the structure, however, it is possible that in the context of blood circulation molecules other than water would be more favorable for this interface.

The purification of native αIIbβ3 from human platelet preserves more glycosylation sites compared to the structures determined from other expression systems. The structural characterization of glycosylation sites in this study offers fundamental building blocks for rigid/homogenous regions of glycans composition and linkage in three-dimensions.^19^ Due to the limitation of naturally purified membrane protein, a mutagenesis study of the glycosylation sites is not applicable to this study. However, these glycosylation sites have already been extensively studied. Due to the flexibility of glycans, the full glycan chain of each site in the αIIbβ3 remains to be determined.

In our study, we solved two co-existing structures in the presence of Mn^2+^, which was proposed to activate the integrin.^29,42,43^ The Mn^2+^ ion is not resolvable due to the resolution limitation, which is common with all structures previously solved in the αIIbβ3 active state.^3,17,31,40^ Instead of an open form of integrin, both forms found in our study adopt the conformation between an inactive and fully active state, which was named the “intermediate” and “pre-active” state. Through comparison of all three structures solved in our study, we hypothesized they could represent a series of states before the active state (Fig. 4).

**Figure 4.**
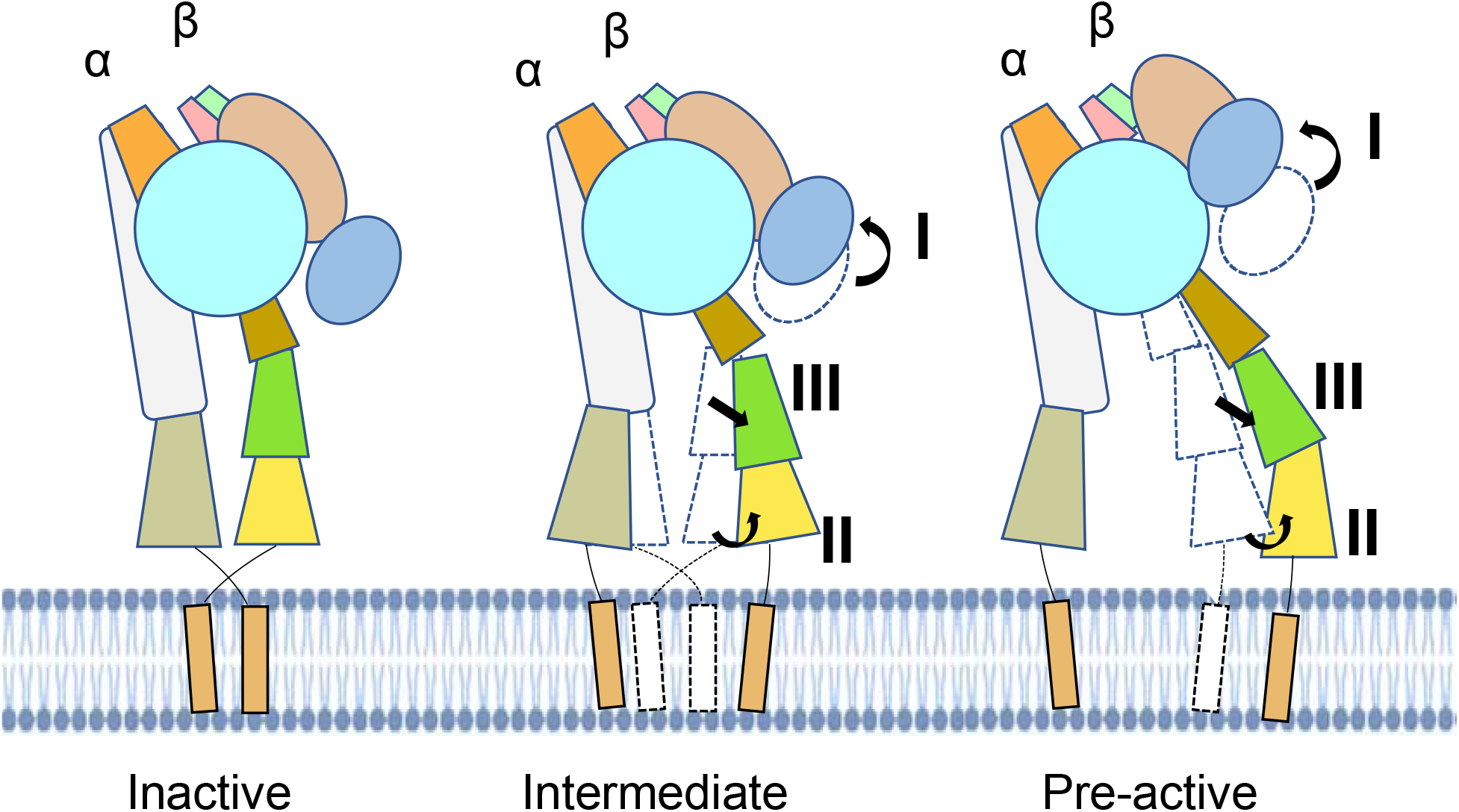
Proposed model for activation transition. The intermediate and pre-active state, both of which show the inter- and intra-molecular separation potential, would exist before the active state adopts a fully open form. Main movements start in the intermediate state at regions I, II, and III, while continuing through the pre-active state. The bent form with separated legs indicates that the leg motion precedes the head lift-up, which explains how the inside out signal transmits.

Upon activation, previous structure of β3-only fragments (contains the headpiece and β-knee regions but lacks the α unit) reported an extended conformation at the I-EGF1/2 junction making the leg almost form a right angle with the head region.^19^ The increased angle between the head and leg regions also exists in our pre-active structure, however, the angle is not as large as it was found in the isolated extended β3. Assuming the leg and head region move away from each other from an acute angle to a straight orientation during activation, it is possible the confirmation found in our study is an initial activation state prior to the proposed activated state. Though the I-EGF1/2 was found to exhibit flexibility to render the conformational change, the rigidity of hybrid/PSI, I-EGF2/3, and I-EGF3/4 is still maintained during activation. This is also consistent with our preactive model, revealing that the interface between I-EGF1/2 is the only changing part during activation (Fig. 3). Specifically, according to another research using anti–HPA-1a alloantibodies to identify the integrin activation, the PSI/I-EGF1 domains were proposed to move away from the I-EGF2 domain during activation.^19^ In this study, by superimposing the head domain of our intermediate and pre-active models, we found I-EGF2 moved closer to the PSI domain, which provides structural evidence to the relative movement between the head and leg regions.

Considering treatment with Mn^2+^ could mimic the inside-out signaling pathway,^17,43–46^ the bent form with separated leg region obtained in our study is in line with the previous hypothetical model,^40^ but provides a substantial and detailed structural basis. Since the actual signal receptor would be the cytoplasmic tails in the inside-out pathway, our structure explains how the leg region moves before and thus induces the head region movement. Therefore, a possible therapeutic design would be more potent if it inhibited the leg movement in the beginning of activation. Since the leg region and head region would be separated as rigid domains, the connection between them, the knee, is presumably a good target site. To be more specific, the interfaces between the αIIb thigh and calf-1 domains and β3 EGF1 and EGF2 domains would exhibit different amino acid profiles and expose different epitopes during the whole activation process, where the inhibitory antibody can intervene and “lock” the integrin in different states accordingly.

## Methods

### Protein extraction and purification

The integrin αIIbβ3 was initially extracted from human platelets obtained from the Gulf Coast Regional Blood Center (Houston, TX, USA). We purified WT full length integrin αIIbβ3 from human platelets by adapting a protocol described previously.^32^ Cells were spun down at 1,200 rpm for 5 min and then resuspended in the CGS buffer for several rounds until red blood cells were fully removed. The final supernatant was collected and centrifuged at 2,500 rpm for 30 min, and the resultant platelet pellet was resuspended using a TBS buffer. After several rounds of homogenization using a Dounce B set, the platelet lysis was spun down at 2000 rpm for 10 min to remove the cell debris followed by another centrifuge at 30,000 rpm for 1 hour to collect the cell membrane fraction. The cell membrane fraction was solubilized at 4°C overnight in the 2% (w/v) Triton X-100 supplemented TBS buffer containing 10 mM Tris-HCl pH 7.4, 150 mM NaCl, 5 mM MgCl2, and 1 mM CaCl_2_. The solubilized membrane protein was collected from the supernatant after centrifugation at 30,000 rpm for 1 hour. The supernatant was then applied to concanavalin-A, an affinity column (Con A Sepharose 4B), and eluted with a buffer containing methyl-α-D-mannopyranoside and 0.01%-0.02% (w/v) Triton X-100. Ion exchange chromatography (MonoQ) and size-exclusion chromatography (Superose 6) were completed using a buffer containing 20 mM Hepes, 150 mM NaCl, DDM (2x-4x CMC), 5 mM MgCl2, and 1 mM CaCl2. Peak fraction was collected for the cryoEM specimen preparation (Supplemental Fig. 1). For integrin αIIbβ3 in the Mn^2+^ condition, the buffer was supplemented with 1 mM MnCl_2_ and 0.2 mM CaCl2 instead of Mg^2+^/Ca^2+^, while using the same purification protocols.

### CryoEM sample preparation and data collection

For cryo-EM sample preparation, a 3 μl aliquot of purified integrin αIIbβ3 was applied onto a 200-mesh R3.5/1 Quantifoil 2nm-Cfilm grid. After applying the sample, the grid was blotted for 3 s and rapidly frozen in liquid ethane using a Vitrobot IV (FEI), with constant temperature and humidity during the process of blotting. The grid was stored in liquid nitrogen before imaging.

Movie stacks were collected at 300 kV on a Krios electron microscope (FEI) with an in-column energy filter (30 eV width) equipped with a direct electron detector K2 Summit camera (Gatan). Images were collected semi automatically by EPU (Thermo Fisher Scientific) in the dose fractionation super-resolution counting mode at a calibrated physical pixel size of 1.07 Å. The images were collected with a defocus range from −1.0 to −2.6 μm. The total exposure time for the dataset was 7 s, leading to a total accumulated dose of 50 electrons Å^2^ on the specimen. Each image stack was fractionated into 35 subframes, each with an accumulation time of 0.2 s per frame. The final frame average was computed from averages of every three consecutive frames to correct beam-induced motion during exposure by MotionCor2.^47^ The image in each frame was weighted according to radiation damage. CTF (Contrast Transfer Function) parameters of the particles in each frame average were determined by the program Patch CTF in cryoSPARC.^48^

In total, 2,758,651 particle images were automatically boxed out by autopicking in cryoSPARC with a box size of 256 × 256 pixels using an averaged sum of 35 raw frames per specimen area. Two-dimensional (2D) reference-free class averages were computed using cryoSPARC. Initial models for every reconstruction were generated from scratch using selected good quality 2D averages with C1 symmetry based on the 2D averaged results. This initial model was low-pass–filtered to 60 Å, and refinements were carried out using cryoSPARC. After several rounds of homogeneous refinement and local refinement, the resolution achieved 3 Å but with a weak signal in the TM region. To improve the quality of the TM region, aligned particles were transferred to RELION^49^ for further processing. To deal with the low signal-to-noise ratio (SNR) and flexibility of the TM region, we employed 3D focus classification to further sort out particles. A spherical mask was created focusing on the TM region and applied to the 3D classification in RELION. The mask was based on TM features and used the detergent shell-reweighted method^50^ to improve TM region SNR. After several rounds of iterative refinement, two helices from αIIb and β3 and the linker between Calf-2 and αIIb helix were resolved, while the connection between β3 tail domain and helix was still elusive due to weak density in the map region. One out of four classes, which appeared as a clear and separated rod-like density, was further processed using the detergent shell denoise and re-weighted method^50^ to improve the SNR (Supplemental Fig. 5). The final map was constructed by the combination of the extracellular domain and TM region. Since within this region, the EM density map is not confidently resolvable for every amino acid due to the flexibility of the leg and TM region, we carried out rigid fitting of structures for the Calf-2 domain from αIIb, the Tail domain from β3, and the TM region. We used previously published TM region structures determined by Nuclear Magnetic Resonance (NMR) (PDB ID: 3FCS for Calf-2 and Tail, 2KNC for TM region) to build these parts of the model, instead of utilizing real-space refinement. The two helices adopted a twisted conformation and went across each other forming a knot at the membrane proximal site. The loop linker between the TM region and whole extracellular domain renders the potential flexibility for the integrin to unwind and separate the helices, which is the prerequisite of activation.

For the map in the Mn^2+^ condition, the cryoSPARC processing workflow is similar to that for native structure (Supplemental Fig. 6). In total, 1,701,301 particles were obtained after iterative 2D classification, and ~10% of particles were used for the initial model reconstruction. Each conformation was obtained from subclasses in the same Mn^2+^ dataset, and corresponding particles were separated for further processing. Two maps showing significant differences were obtained with the particle ratio 1:1.02. For state **III**, the EM map was refined to a proper resolution where domain information could be seen, while details for the secondary structure were still missing (Fig. 3). Individual domain from integrin αIIbβ3 was rigid-docking in the density. After individual refinement for each particle set, the two maps were finally refined to 3.09 Å and 5.46 Å respectfully.

### Cryo-EM model building and refinement

We started de novo model building for full-length inactive integrin αIIbβ3 by docking the extracellular domain (PDBID:3FCS)^15^ and TM region (PDBID:2KNC).^51^ The model was refined against the corresponding map using PHENIX^52^ in real space with the secondary structure and geometry restraints. Since the EM density map is not confidently resolvable for every amino acid at the leg and TM regions due to the flexibility, we carried out rigid fitting of structures of the Calf-2 domain from αIIb, the Tail domain from β3, and the TM region. We used previously published TM region structures determined by Nuclear Magnetic Resonance (NMR) (PDB ID 3FCS for Calf-2 and Tail, 2KNC for TM region) to build these parts of the model, instead of utilizing real-space refinement. The model was then manually refined and adjusted in Coot.^53^ The model quality was validated using PHENIX and the local resolution was estimated using cryoSPARC. The aforementioned workflow applied to structure refinement in the inactive form and intermediate form, while the pre-active form was only applied with the rigid body-fitting for each domain due to low resolution. Detailed statistics for model building and refinements are given in table (S1). The final maps and models were submitted to the Electron Microscopy Data Bank (EMDB) (accession no) and (accession no) and PDB (accession no).

## Author Contributions

Z.W. conceived and supervised the project. M.S. guided T.H., H.W., and Z.M. in preparing the integrin sample. T.H. and H.W. performed the data collection, processing, and analysis. T.H. made the movies. T.H., M.S., H.W., and Z.W. wrote the manuscript with other authors’ input.

## Acknowledgement

We thank Guizhen Fan, Xinzhe Yu, Joshua I Rosario Sepulveda, Steve Ludtke and Theodore Wensel for suggestions and comments on the experiment and manuscript. This work is supported by (R01GM143380 and R01HL162842) BCM BMB department seed funds to Z.W. CryoEM data was collected at the Baylor College of Medicine CryoEM ATC, which includes equipment purchased under support of CPRIT Core Facility Award RP190602.

---

**Supplemental Figure 1.**
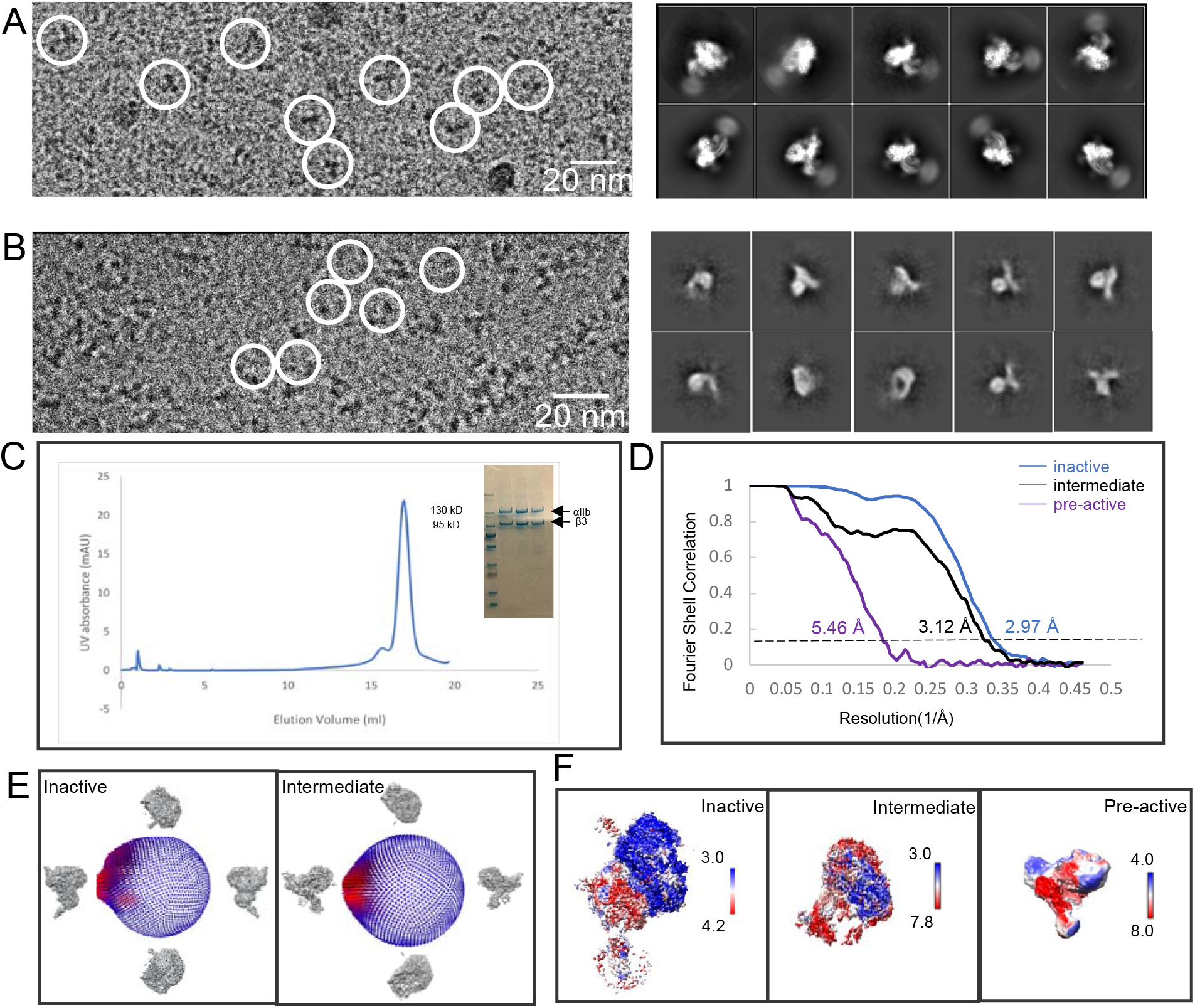
The sample and SPA of integrin in the inactive state and intermediate state. A) Representative micrograph and 2d classification of integrin in inactive state. B) Representative micrograph and 2d classification of integrin in intermediate state. C) SEC-FPLC profile D) The Fourier shell correlation (FSC) curves of the three reconstructions using the gold-standard criteria (FSC=0.143). E) The angular distribution plot for two maps. F) The local resolution variations in the three cryo-EM maps.

**Supplemental Figure 2.**
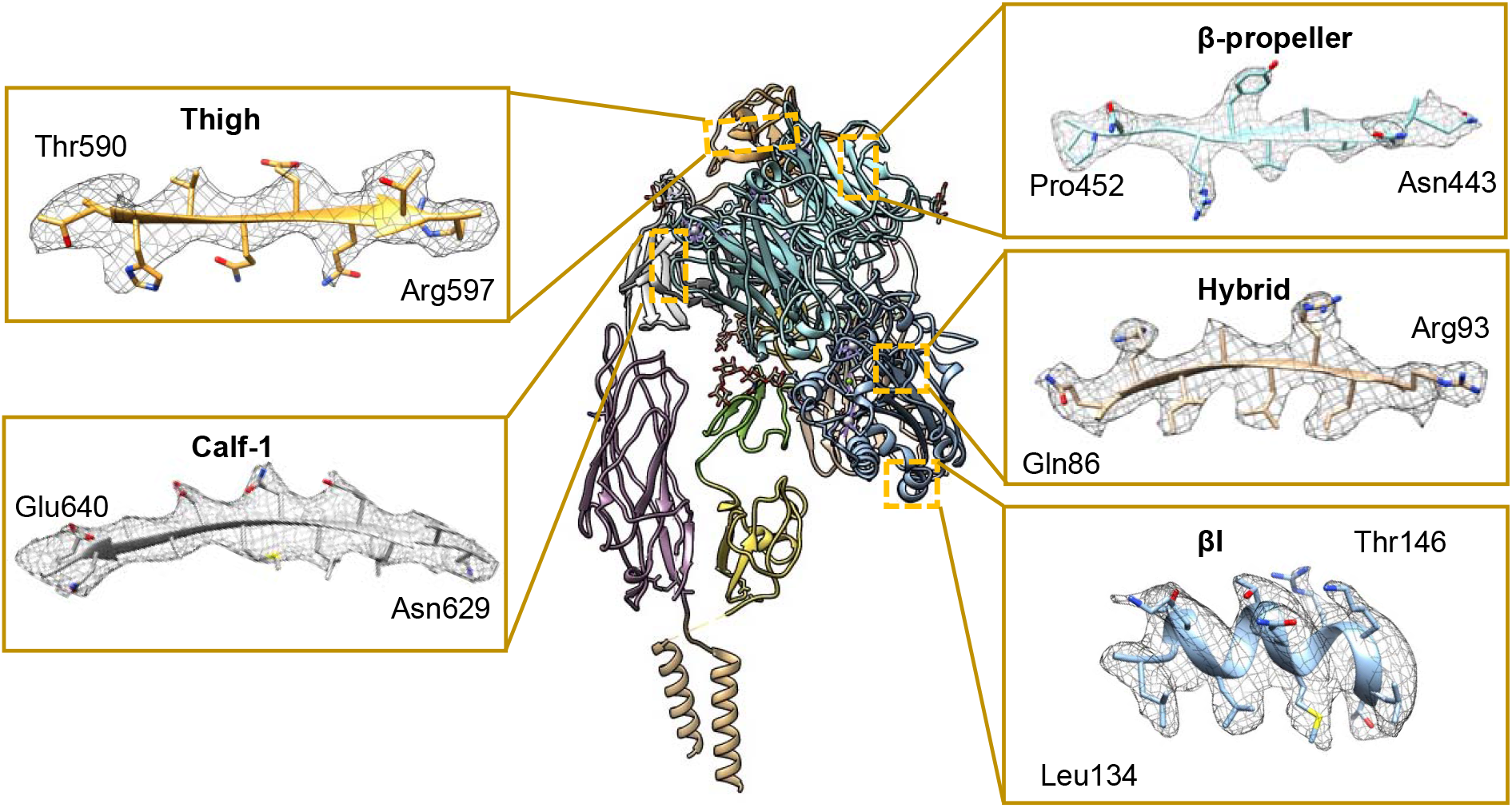
Representative sample of the 2.9 Å inactive cryo-EM map of αIIβ3 integrin showing side-chain densities along with the modeled structure.

**Supplemental Figure 3.**
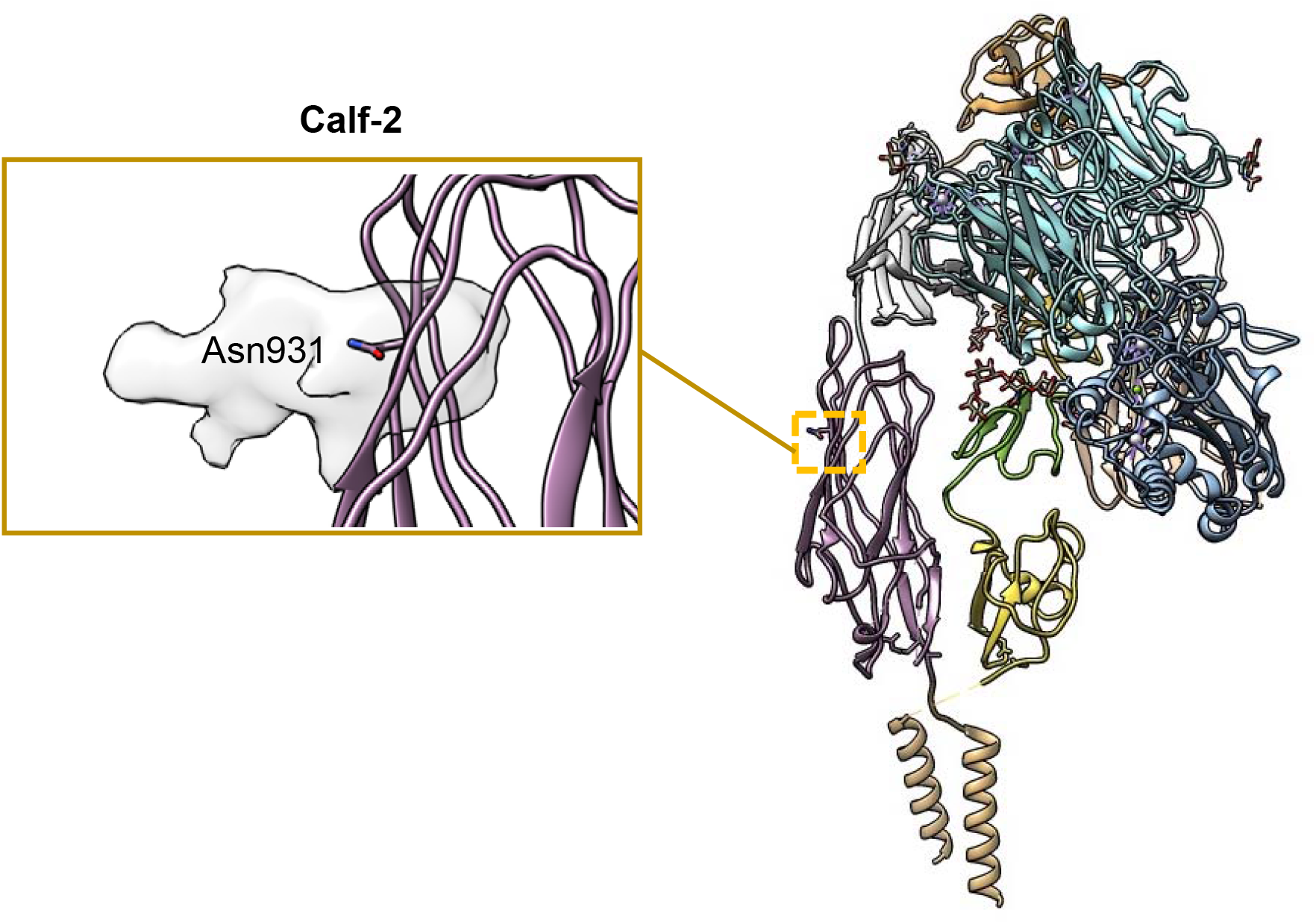
Extra density of the glycosylation for Asn931 is showed in the low-pass filter map (resolution ≈ 7.5 Å).

**Supplemental Figure 4.**
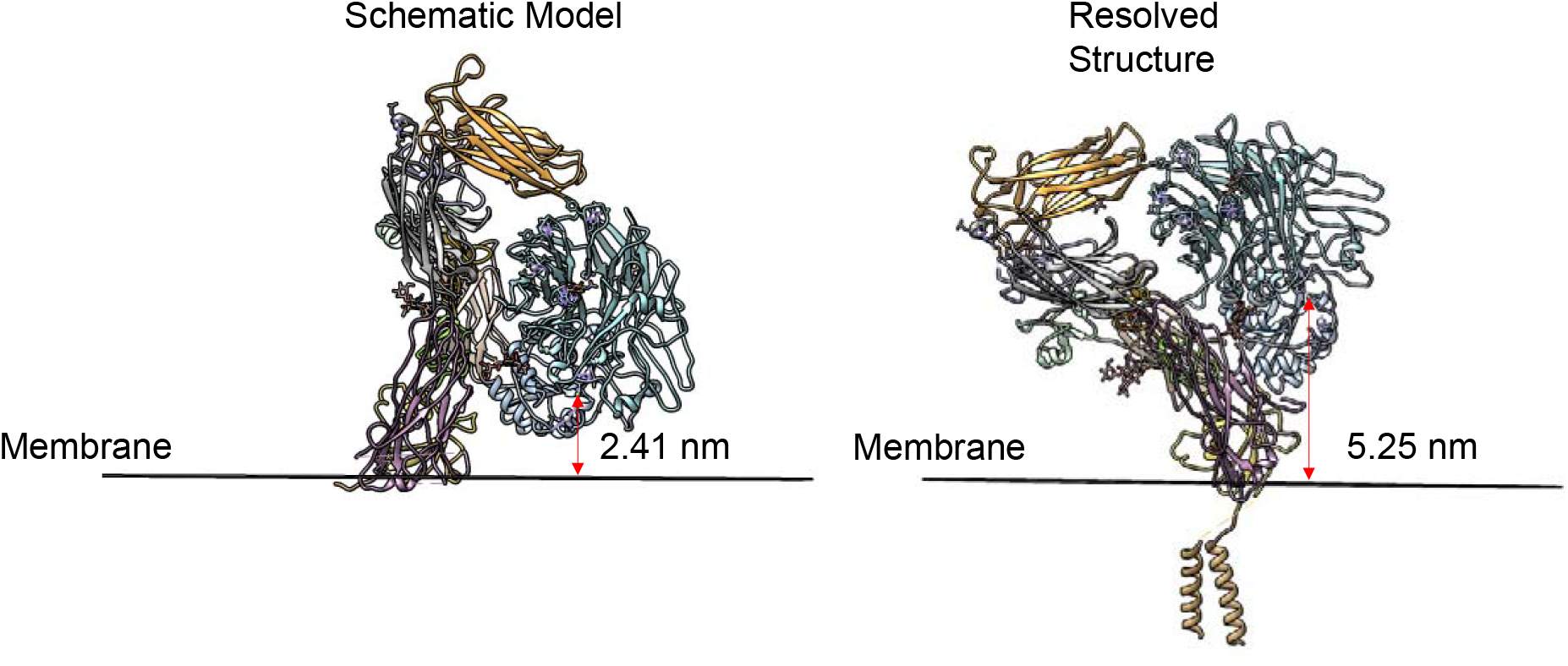
Distance between MIDAS (use the coordinate of Mg^2+^) and cell membrane. In the previous schematic model, cell membrane plane is perpendicular to “leg” region. The distance is 2.41 nm (left). In our model, cell membrane plane is perpendicular to TM region and extracellular domain is tilted compared to previous model. The distance is 5.25 nm(right).

**Supplemental Figure 5.**
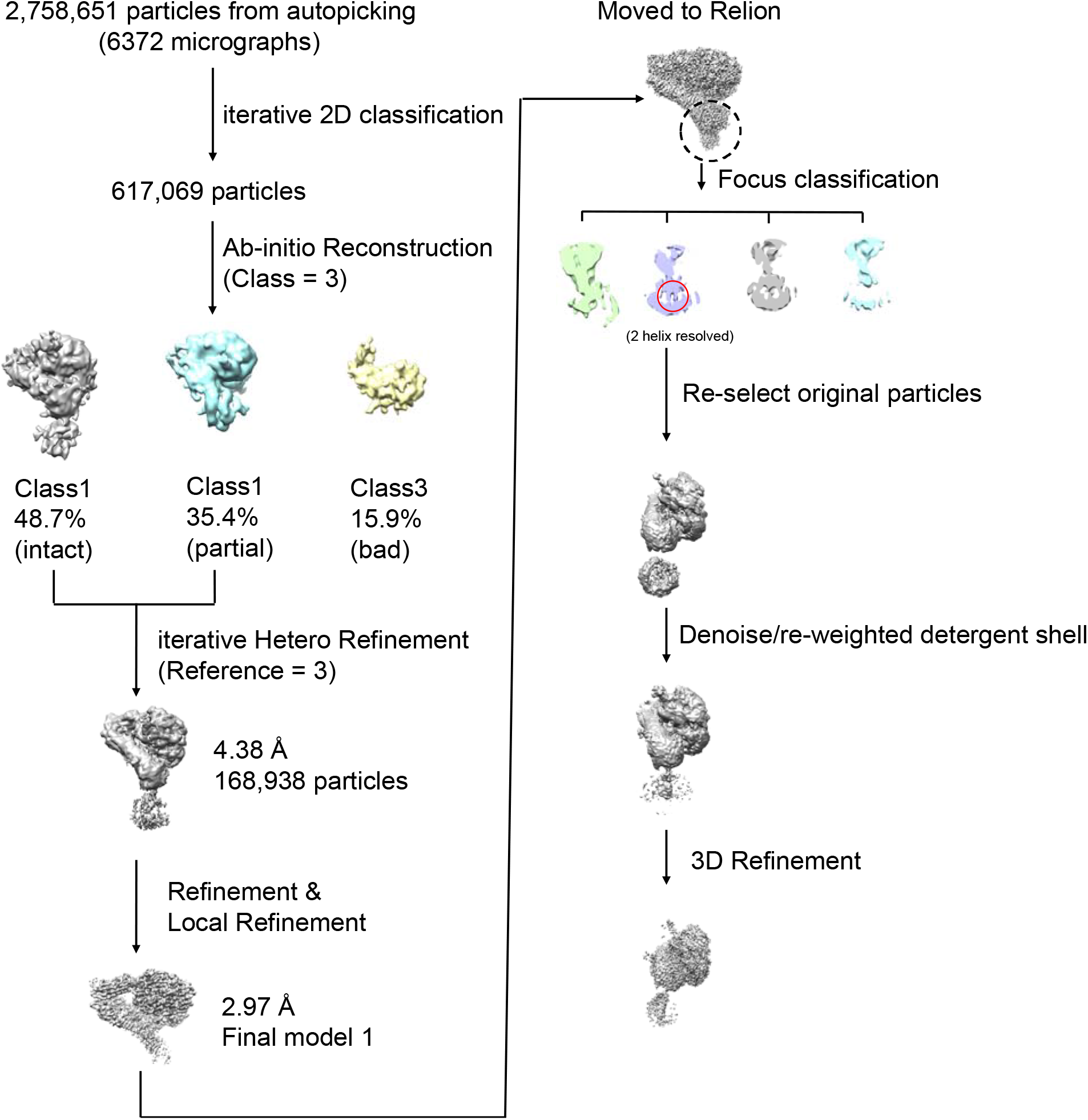
CryoEM data processing for αIIβ3 in Ca^2+^ and Mg^2+^ condition.

**Supplemental Figure 6.**
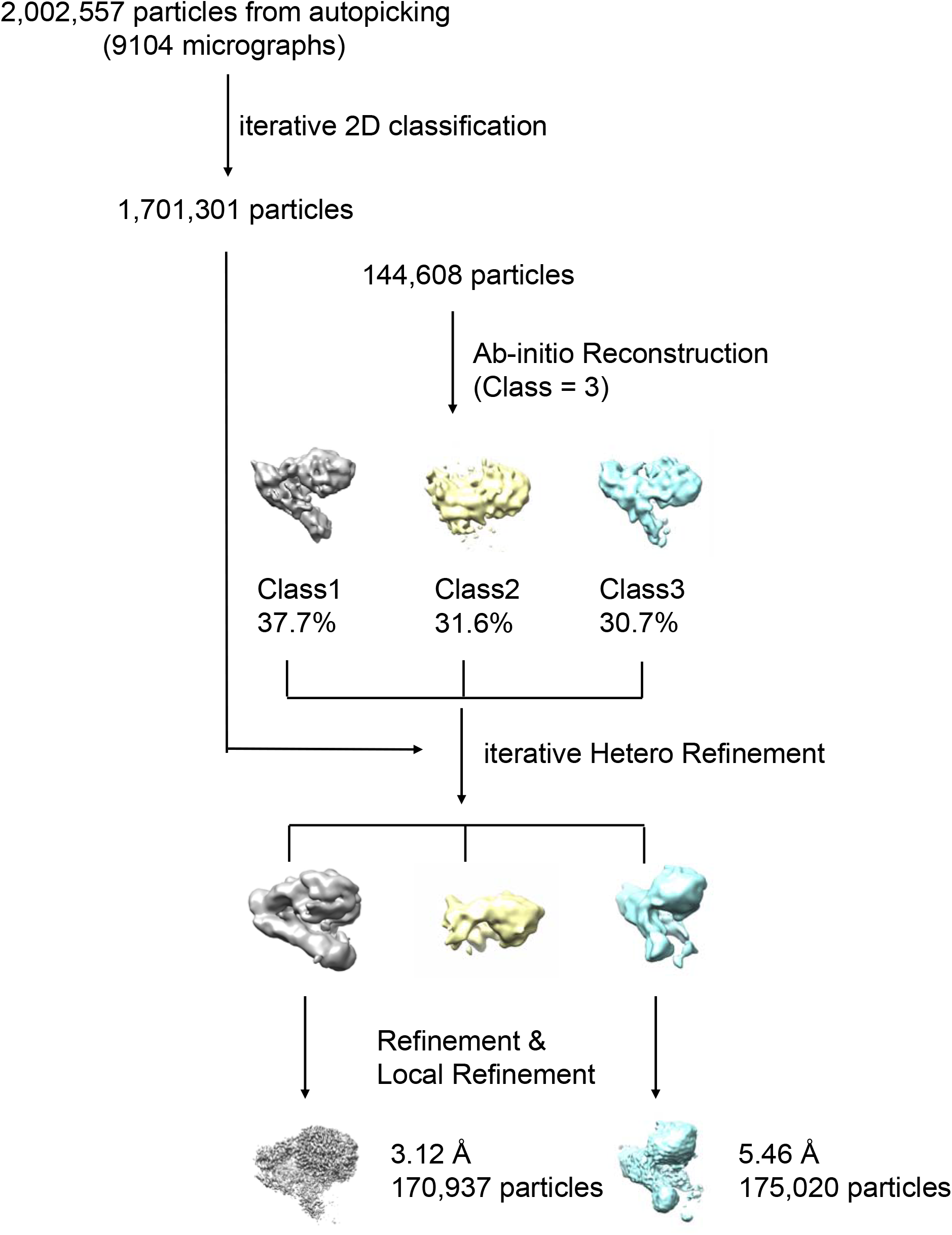
CryoEM data processing for αIIβ3 in Ca^2+^ and Mn^2+^ condition.

**Supplemental Movie 1.** Shown are the cryoEM density maps and structure for the inactive state of αIIβ3 in Ca^2+^ and Mg^2+^.

**Supplemental Movie 2.** Shown are the cryoEM density maps and structure for the intermediate state of αIIβ3 in Ca^2+^ and Mn^2+^.

**Supplemental Movie 3.** Transition morph movie from inactive state to intermediate state and preactive state of αIIβ3.

**Supplemental Table 1.** Cryo-EM data collection, refinement and validation statistics.

| | Inactive αIIbβ3<br>(EMDB-XXXX)<br>(PDB-XXXX) | Intermediate αIIbβ3<br>(EMDB-XXXX)<br>(PDB-XXXX) | (EMDB-XXXX) |
| --- | --- | --- | --- |
| **Data collection and processing** |  |  |  |
| Instrument | Krios (FEI) | Krios (FEI) | Krios (FEI) |
| Director | K2 | K2 | K2 |
| Magnification | 130K | 130K | 130K |
| Voltage (kV) | 300 | 300 | 300 |
| Electron dose (e-/Å<sup>2</sup>) | 1.43 | 1.43 | 1.43 |
| Defocus range (μm) | -1.0 to -2.6 | -1.0 to -2.6 | -1.0 to -2.6 |
| Pixel size (Å) | 1.07 | 1.07 | 1.07 |
| Symmetry imposed | C1 | C1 | C1 |
| Micrograph collected (N) | 6372 | 9104 | 9104 |
| Initial particle images(N) | 2,758,651 | 2,002,557 | 2,002,557 |
| Final particle images(N) | 168,938 | 170,937 | 175,020 |
| Map resolution (Å) | 2.97 | 3.12 | 5.46 |
| FSC threshold | 0.143 | 0.143 | 0.143 |
| **Refinement** |  |  |  |
| Initial model used(PDBs) | 3FCS | 3FCS |  |
| Model resolution(Å) | 3.49 | 3.90 |  |
| FSC threshold | 0.5 | 0.5 |  |
| **Validation** |  |  |  |
| B factors(Å<sup>2</sup>)(mean) | 61.79 | 49.91 |  |
| R.m.s. deviations |  |  |  |
| Bond lengths (Å) | 0.007 | 0.005 |  |
| Bond angles (°) | 1.412 | 0.777 |  |
| MolProbity score | 1.96 | 2.27 |  |
| Clash score | 10.66 | 18.00 |  |
| Poor rotamers (%) | 0.53 | 0.37 |  |
| **Ramachandran plot** |  |  |  |
| Favoured (%) | 93.74 | 91.06 |  |
| Allowed (%) | 6.11 | 8.75 |  |
| Disallowed (%) | 0.15 | 0.19 |  |

